# Astrocyte subtype-specific expression of the sodium-coupled citrate transporter SLC13A5 and citrate metabolism genes across Alzheimer’s disease pseudoprogression: a single-nucleus RNA sequencing analysis of the human middle temporal gyrus

**DOI:** 10.64898/2026.06.05.730472

**Authors:** Patrícia Fernanda Schuck, Gustavo da Costa Ferreira, Hércules Rezende Freitas

## Abstract

The sodium-coupled citrate transporter NaCT (SLC13A5) imports extracellular citrate into cells. In the CNS, SLC13A5 is described to be expressed predominantly in neurons. Cytosolic citrate levels rely on citrate generated in mitochondria and imported from other CNS cells, regulating intermediary metabolism and supplying acetyl-CoA for lipid synthesis and histone acetylation. Despite evidence for NaCT’s role in neurometabolic homeostasis, its transcriptional behavior across Alzheimer’s disease (AD) progression and across astrocyte subtypes remains uncharacterized at single-cell resolution. We analyzed single-nucleus RNA sequencing data from 1,378,211 nuclei across 84 donors in the Seattle Alzheimer’s Disease Brain Cell Atlas (SEA-AD) Middle Temporal Gyrus dataset to profile SLC13A5 and seven citrate metabolism genes across a continuous AD pseudoprogression score. SLC13A5 expression was restricted to astrocytes (∼20% prevalence) and concentrated in the Astro 2 supertype (24.0%), a homeostatic subtype characterized by low C3 (1.6%) and CD44 (5.5%), which expanded with pseudoprogression (Spearman rho = +0.345, FDR < 0.001). The A1-reactive Astro 3 supertype, where SLC13A5 prevalence was 0.87%, declined concordantly (rho = -0.393). Opposing compositional and transcriptional forces produced apparent stability in overall SLC13A5 prevalence. SLC13A3 and ACO1 showed progressive donor-level declines correlating with Braak stage and Thal phase (rho range: -0.307 to -0.349, FDR < 0.01). APOE4 carriers exhibited lower SLC13A5 prevalence specifically within Astro 2 nuclei (median 17.6% vs. 25.9%; Wilcoxon p = 0.025), though this association did not survive multivariate regression. No difference in Astro 2 SLC13A5 expression was detected between cognitively resilient and expected-AD donors with equivalent high Braak burden (p = 0.888). Contrary to the prevailing description of NaCT as a neuronal transporter, SLC13A5 expression in the SEA-AD MTG dataset is restricted to astrocytes, concentrated in the homeostatic Astro 2 subtype, and maintained as this subtype expands with advancing AD pathology. Supertype-resolved SLC13A5 and SLC13A3 expression provide more informative readouts of astrocytic metabolic state than bulk measurements.

## 1. Introduction

Citrate occupies a central node in neurometabolic homeostasis. As a TCA cycle intermediate exported from astrocytic mitochondria via the SLC25A1 carrier, citrate reaches the cytoplasm and the extracellular space. Citrate can be taken up from extracellular milieu by the sodium-coupled transporter NaCT, encoded by *SLC13A5* [1]. Within astrocytes, cytoplasmic citrate is cleaved by ATP-citrate lyase (ACLY) to generate acetyl-CoA, the obligate substrate for histone acetylation and lipid synthesis [2]. Among the possible contributors to this acetyl-CoA pool, astrocytic citrate import via NaCT is one candidate source; however, the relative contribution of extracellular citrate versus mitochondrially derived citrate to cytoplasmic acetyl-CoA for histone acetylation and lipid synthesis remains to be established. Loss-of-function mutations in *SLC13A5* cause neonatal epileptic encephalopathy with impaired brain development in humans [3,4], confirming that citrate transport is essential for neural homeostasis from early life onward. In *Drosophila*, loss of the INDY/*SLC13A5* ortholog extends lifespan [5], and mammalian *SLC13A5* knockout reduces adipogenesis through altered mitochondrial biogenesis and energy sensing in peripheral metabolic tissues [6], suggesting that NaCT-mediated citrate flux connects to energy sensing and lipid metabolism well beyond simple carbon shuttling.

In Alzheimer’s disease (AD), astrocyte metabolism is broadly disrupted [7,8]. Reactive astrogliosis, impaired glutamate uptake, dysregulated lipid metabolism, and mitochondrial dysfunction have all been documented in post-mortem tissue and in single-nucleus transcriptomic studies [9,10]. Astrocytes are not a monolithic population: they encompass homeostatic and reactive subtypes with distinct gene expression programs, spatial distributions, and disease-associated trajectories [7,9]. The A1/A2 classification, now widely recognized as an oversimplification of a continuous reactive spectrum [11,12], identified complement component *C3* and *CD44* as markers of neurotoxic reactive astrocytes induced by microglial pro-inflammatory signals (IL-1*α*, TNF, C1q) [11]. Weighted gene co-expression network analysis of large-scale bulk brain transcriptomics has begun to characterize biological processes associated with *SLC13A5* expression across the human lifespan [13], but whether *SLC13A5*-expressing astrocytes belong to homeostatic or reactive subpopulations, and how that membership shifts across AD neuropathological progression, has never been resolved at the single-cell level. Furthermore, NaCT is canonically described as a sodium-coupled citrate transporter expressed predominantly in neurons [1], yet whether this neuronal expression pattern holds across all brain regions, has not been systematically examined. The possibility that SLC13A5 expression is concentrated in astrocytes rather than neurons in specific cortical regions, and that this distribution shifts with neuropathological burden, represents an unresolved question with direct implications for understanding how citrate metabolism is regulated at the glial-neuronal interface in the aging and AD brain.

The Seattle Alzheimer’s Disease Brain Cell Atlas (SEA-AD) offers the scale needed to address this question directly [14]. Its Middle Temporal Gyrus dataset spans 1,378,211 nuclei from 84 donors with varying neuropathological burden and includes a continuous per-cell pseudoprogression score reflecting aggregate AD-associated transcriptional change independent of clinical diagnosis. This score makes it possible to track gene expression across a disease continuum rather than forcing donors into binary groups, which is more sensitive to graded transcriptional shifts than neuropathological staging categories alone. Here, building on prior bulk transcriptomic characterization of brain *SLC13A5* coexpression networks [13], we provide the first cell-type-resolved, single-nucleus characterization of *SLC13A5* expression and seven additional citrate metabolism genes (*SLC13A3, SLC25A1, ACLY, ACO1, ACO2, IDH1, IDH2*) across 24 cell subclasses and six astrocyte supertypes in the SEA-AD MTG dataset. We ask whether NaCT expression changes with pseudoprogression, which astrocyte subtype carries the signal, and whether clinical and genetic variables (Braak stage [15], Thal phase [16], CERAD score [17], APOE4 genotype) modulate expression of this transporter. The results reveal that *SLC13A5* expression in the MTG is unexpectedly restricted to astrocytes, with near-absent detection in neuronal subclasses, and that apparent stability in overall astrocyte-level prevalence across pseudo-progression conceals a subtype-level reorganization of the astrocytic compartment, with specific implications for how citrate metabolism is maintained, and may be compromised, as AD pathology accumulates.

## 2. Materials and methods

### 2.1. Dataset

We used single-nucleus RNA sequencing (snRNA-seq) data from the Seattle Alzheimer’s Disease Brain Cell Atlas (SEA-AD) Middle Temporal Gyrus release (version 2024-02-13) [14], available via the Allen Brain Cell Atlas (https://portal.brain-map.org/atlases-and-data/rnaseq/human-mtg-10x_sea-ad). The dataset comprises 1,378,211 nuclei from 84 human donors (42 with dementia, 42 without dementia), processed with the 10x Genomics Chromium 3’ v3 platform. A continuous pseudoprogression score (0–1) provided by the SEA-AD consortium reflects the aggregate burden of AD-associated neuropathological changes inferred from transcriptomic profiles; cells without a valid pseudoprogression score were excluded from trajectory analyses. Donor-level metadata extracted from the h5ad file included Braak stage (I–VI) [15], CERAD score (1–4) [17], Thal phase (0–5) [16], APOE genotype, cognitive status (No dementia/Dementia/Reference), age at death, and post-mortem interval (PMI).

### 2.2. Cell type annotation

Cell subclass and supertype annotations were taken directly from the SEA-AD taxonomy, which assigns each nucleus to one of 24 subclasses and 139 supertypes using hierarchical reference-mapping. For astrocyte-specific analyses, we used the six astrocyte supertypes: Astro 1 through Astro 6. Nuclei annotated as “Reference” donors (neurotypical reference cases without neuropathological burden) were present in the full dataset but underrepresented in astrocyte analyses (84 donors had valid pseudoprogression scores). All 24 cell subclasses, including glutamatergic and GABAergic neuronal populations, were retained for the full-dataset dot plot analysis (Figure 1E), which characterizes the cell-type specificity of each gene’s expression and provides the empirical basis for the near-absent SLC13A5 detection observed in neuronal subclasses. Supertype-resolved trajectory, neuropathological, and genetic analyses were restricted to astrocyte nuclei (i.e., single-nucleus RNA-seq capture units from cells assigned to the Astrocyte subclass) given the near-zero neuronal prevalence, which precludes meaningful subtype-level quantification in that compartment.

**Figure 1.**
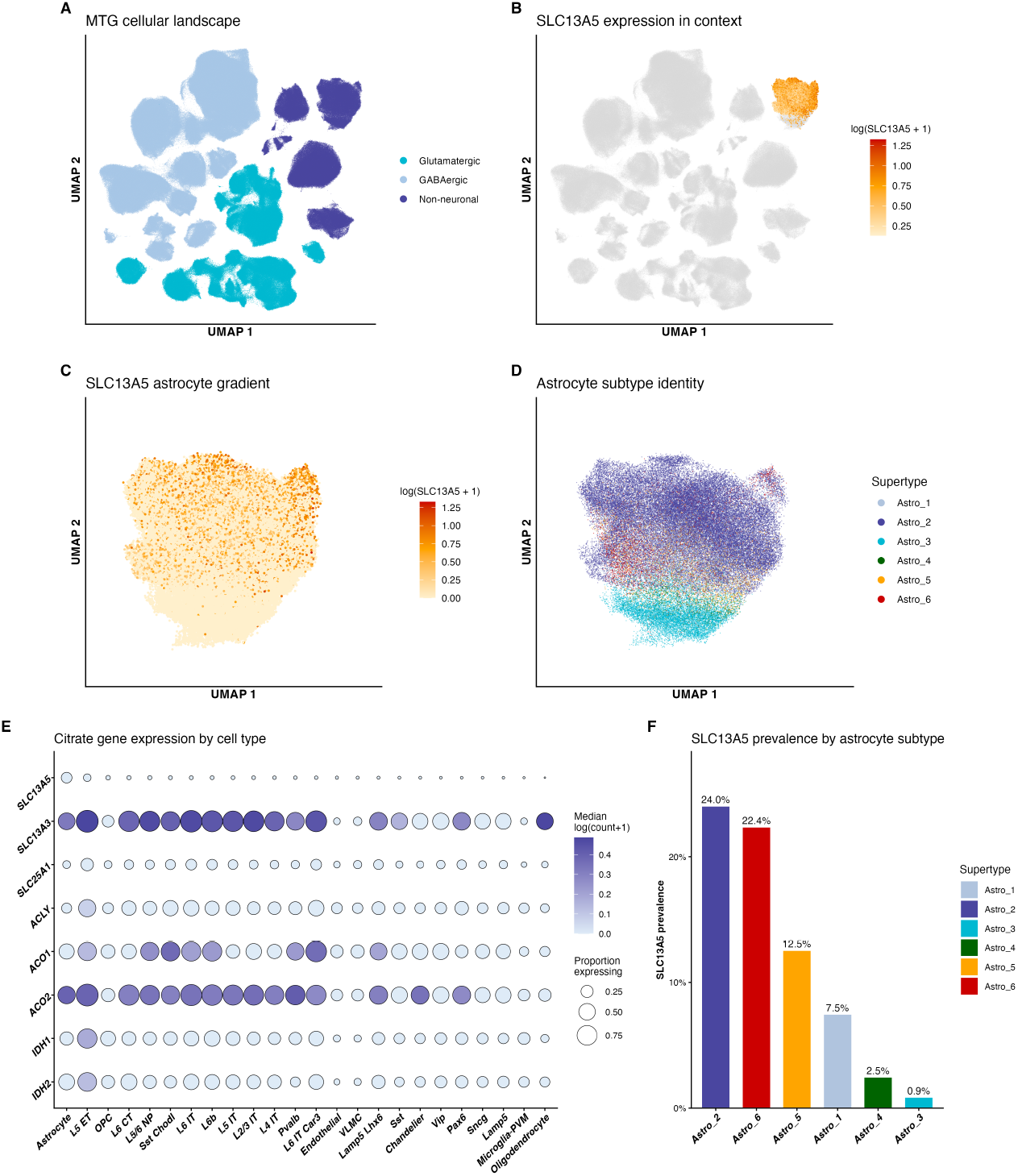
SLC13A5 expression landscape in the SEA-AD middle temporal gyrus. (A) UMAP of 1,378,211 nuclei by broad cell class. (B) SLC13A5-expressing astrocytes (orange-red) overlaid on the full UMAP. (C) SLC13A5 expression within the astrocyte cluster (zoomed). (D) Astrocyte supertype identity in the same zoomed view; Astro 2 is the primary SLC13A5-expressing population. (E) Proportion expressing (dot size) and median log-normalized count (dot color) for eight citrate metabolism genes across 24 cell subclasses. (F) SLC13A5 prevalence across the six astrocyte supertypes.

It should also be noted that Reference donors, neurotypical individuals without significant neuropathological burden, are underrepresented in this dataset relative to donors across the AD spectrum. Whether SLC13A5 expression is detectable in neuronal subclasses in the neurotypical MTG, and whether its apparent absence in neurons reflects a disease-associated transcriptional change rather than a constitutive regional pattern, could not be fully evaluated with the current donor composition. This limitation should be considered when interpreting the cell-type specificity of SLC13A5 expression reported here.

### 2.3. Gene expression extraction

Normalized log-transformed expression values were extracted from the X matrix of the h5ad file using chunk-wise reading via the `rhdf5` package in R [18], avoiding loading the full 1.37M *×* 36,601 matrix into memory. Values in X represent CP10K-normalized, log1p-transformed expression (i.e., log(count *×* 10,000/library size + 1)). The indptr array was read with 64-bit float conversion (`bit64conversion = “double”)` to handle cumulative non-zero counts exceeding the 32-bit integer limit. For all binary detection analyses (proportion of nuclei with normalized value > 0), the CP10K + log1p transformation is monotone, so the detection threshold is equivalent to *≥*1 raw unique molecular identifier (UMI) per nucleus. Genes analyzed: *SLC13A5, SLC13A3, SLC25A1, ACLY, ACO1, ACO2, IDH1, IDH2* (citrate metabolism panel); *GFAP, VIM, C3, ALDH1L1, AQP4, S100B, CD44, LCN2* (reactive astrocyte markers).

### 2.4. Expression metrics

For each gene, two complementary metrics were computed: (1) prevalence (binary margin): the proportion of nuclei with count > 0, reflecting the fraction of cells expressing the gene; and (2) intensity (continuous margin): mean log(count + 1) among expressing nuclei only, reflecting expression level conditional on detection. Unless otherwise stated, analyses use prevalence as the primary outcome, consistent with the extensive margin interpretation appropriate for sparse snRNA-seq data.

To evaluate whether *SLC13A5* detection in non-astrocyte subclasses reflected genuine expression or sequencing-depth-driven stochastic capture, cells were stratified by total UMI count quartile within each subclass and prevalence was computed per quartile. Astrocyte prevalence showed modest depth dependence (Q1: 10.3% at median 4,054 UMIs per cell; Q4: 29.8% at 19,950 UMIs), consistent with a genuinely expressed gene maintaining a robust detection floor at low sequencing depth. L5 ET neurons showed strong depth dependence (Q1: 2.6% at median 21,074 UMIs; Q4: 13.8% at 133,551 UMIs). Critically, astrocyte Q1 cells with five-fold lower median library sizes than L5 ET Q1 cells still showed four-fold higher prevalence (10.3% vs. 2.6%), indicating that the L5 ET signal is predominantly attributable to sequencing depth rather than constitutive expression. Total UMI counts per cell were obtained from the `Number of UMis` field of the h5ad file.

### 2.5. Pseudoprogression trajectory analysis

Astrocyte nuclei with valid pseudoprogression scores (n = 67,419) were binned into 20 equal-width bins. Per-bin prevalence and mean intensity were computed for each gene. LOESS smoothing (span = 0.75) was applied to visualize trajectories. Spearman rank correlations were computed between pseudoprogression score and binary expression status for each gene across all 67,419 astrocyte nuclei (cell-level analysis), and separately across 84 donor-level aggregates using mean pseudoprogression score per donor as the independent variable. Fisher z-transformation was used to compute 95% confidence intervals for Spearman rho. Segmented regression (R package `segmented` [19]) was applied to the binned prevalence data to test for a change-point in the *SLC13A5* trajectory. Logistic regression modeled the binary *SLC13A5* outcome (expressed vs. not) as a function of pseudoprogression score and sex. All p-values were adjusted using the Benjamini-Hochberg (BH) false discovery rate procedure within each analysis block.

### 2.6. Astrocyte supertype composition analysis

Donor-level proportions of each of the six astrocyte supertypes were computed as the fraction of that donor’s astrocyte nuclei assigned to each supertype. Spearman correlations between these proportions and donor mean pseudoprogression score were computed. Reactive astrocyte markers (*GFAP, VIM, C3, ALDH1L1, AQP4, S100B, CD44, LCN2*) were extracted for astrocyte nuclei and compared between Astro 2 and Astro 3 using Wilcoxon rank-sum tests with BH correction.

### 2.7. Co-expression analysis

Phi coefficients (binary Pearson correlation equivalents) were computed between all pairs of the eight citrate metabolism genes across the binary expression matrix (67,419 astrocyte nuclei *×* 8 genes). Integer overflow was prevented by wrapping `sum()` calls in `as.numeric()` before computing the denominator product. Conditional prevalence P(gene B expressed | gene A expressed) was computed for all gene pairs. *SLC13A5*-specific enrichment ratios compared co-expression prevalence in *SLC13A5*+ vs. *SLC13A5*-nuclei.

### 2.8. Neuropathological and genetic associations

Donor-level Spearman correlations between gene prevalences and Braak stage [15], Thal phase [16], CERAD score [17], age at death, and PMI were computed using the 84-donor aggregate dataset. APOE4 status was dichotomized: APOE4+ (genotypes E3/E4, E4/E4) vs. APOE4-(E2/E2, E2/E3, E2/E4, E3/E3) [20]. E2/E4 carriers were assigned to the APOE4-group because the protective APOE2 allele substantially attenuates APOE4-associated AD risk, making this genotype biologically ambiguous for a binary APOE4 contrast. Wilcoxon rank-sum tests compared gene prevalence between APOE4 groups, between cognitive status groups (No dementia vs.∼Dementia), and between cognitive resilience groups (Resilient: High Braak + No dementia; Expected AD: High Braak + Dementia; Low pathology: Low Braak) [21]. Multivariate OLS regression modeled logit-transformed *SLC13A5* prevalence in Astro 2 nuclei as a function of standardized pseudoprogression, age, APOE4 status, and sex across 84 donors. Variance inflation factors confirmed absence of multicollinearity (all VIF < 1.3).

### 2.9. Statistical software

All analyses were performed in R (version *≥* 4.3) [22]. Key packages used: `tidyverse` [23], `Matrix` [24], `broom` [25], `car` [26], `patchwork` [27], `ggplot2` [28], `scales` [29]. A fully reproducible code is available in the supplementary companion statistical report [30].

## 3. Results

### 3.1. SLC13A5 expression is restricted to astrocytes and concentrated in the Astro 2 supertype

Among 1,378,211 nuclei across 24 cell subclasses in the SEA-AD MTG dataset, *SLC13A5* was expressed almost exclusively in astrocytes. The full cellular landscape of the SEA-AD dataset and the spatial distribution of *SLC13A5*-expressing cells are shown in Figure 1A-D. Approximately 20% of astrocyte nuclei were *SLC13A5*-positive. This prevalence was robust across sequencing depth, remaining above 10% even in the lowest library-size quartile of astrocyte nuclei (median 4,054 UMIs per cell; Figure 1E). L5 ET neurons showed 7.7% overall prevalence but strong depth dependence (2.6% in the lowest quartile at median 21,074 UMIs; 13.8% in the highest quartile at 133,551 UMIs), consistent with stochastic capture of rare transcripts in deeply sequenced cells rather than constitutive expression. At library sizes equivalent to typical astrocytes (*∼*4,000–10,000 UMIs), SLC13A5 detection in L5 ET neurons would approach background levels. All remaining 22 subclasses showed prevalence below 2.1% regardless of sequencing depth (Figure 1E). This cell-type specificity, visible across all 24 subclasses in the dot plot, contrasts with the canonical description of NaCT as a neuronal transporter [1] and is a primary finding of this study. Among the six transcriptomically defined astrocyte supertypes, *SLC13A5* expression was concentrated in Astro 2 (24.0% prevalence) and nearly absent from Astro 3 (0.87%; Figure 1F). The three other supertypes with sufficient nuclei for analysis (Astro 1, Astro 5, Astro 6) showed intermediate prevalence values ranging from approximately 8% to 15%.

*SLC13A3*, encoding the sodium-dependent dicarboxylate transporter NaDC3/SDCT2 [31], showed astrocyte-enriched expression at markedly higher prevalence (*∼*57%) and was additionally detected across several non-astrocyte subclasses.

Of the citrate metabolism enzymes, *ACO1, ACO2*, and *IDH1*/*IDH2* showed broad expression across multiple subclasses, while *SLC25A1* and *ACLY* showed enrichment in both astrocytes and neurons, at lower overall prevalence than *SLC13A3* (Figure 1E).

### 3.2. Overall SLC13A5 prevalence is stable across pseudoprogression

Cell-level Spearman correlation between pseudoprogression score and binary *SLC13A5* expression across 67,419 astrocyte nuclei yielded rho = -0.008 (FDR = 0.061; Figure 2C), indicating no meaningful monotonic trend. This null result was confirmed by logistic regression (pseudoprogression score OR not significant), by supertype-stratified analyses showing heterogeneous and partially opposing directional signals across supertypes, and by segmented regression identifying a weak non-monotonic shape with a potential inflection near pseudoprogression score 0.52 (SE = 0.10). At the donor level (n = 84), *SLC13A5* prevalence did not significantly correlate with mean pseudoprogression score (FDR > 0.05; Figure 2D).

**Figure 2.**
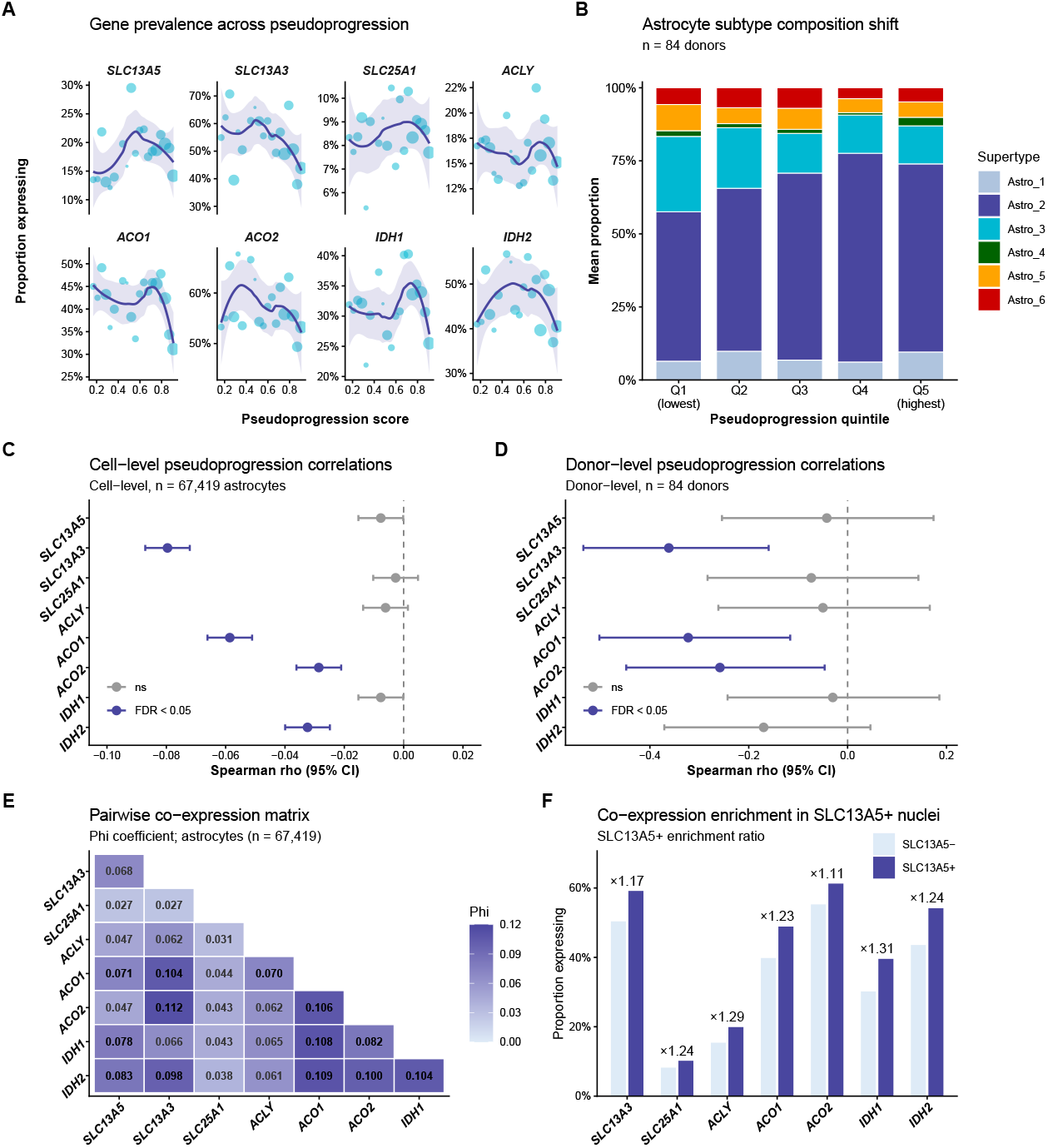
Pseudoprogression trajectories and co-expression architecture of citrate metabolism genes. (A) LOESS-smoothed prevalence trajectories for eight genes across pseudoprogression (20 bins; shading = 95% CI). (B) Astrocyte supertype composition per pseudoprogression quintile (n = 84 donors); Astro 2 expands and Astro 3 contracts. (C) Cell-level Spearman correlations between pseudoprogression score and binary gene expression (n = 67,419 astrocyte nuclei). (D) Donor-level Spearman correlations using aggregated prevalence (n = 84 donors). (E) Phi coefficient matrix for pairwise binary co-expression. (F) Proportion of each gene expressed in SLC13A5+ versus SLC13A5-nuclei; enrichment ratios shown above SLC13A5+ bars.

In contrast, *SLC13A3, ACO1, IDH2*, and *ACO2* all showed statistically significant negative cell-level correlations with pseudoprogression (FDR < 0.001; Figure 2A), with *SLC13A3* showing the largest effect (rho = -0.080). At the donor level, *SLC13A3* exhibited a substantial negative correlation (rho = -0.362, FDR < 0.05), confirming that the cell-level signal represents genuine inter-donor variability rather than pseudoreplication artifact.

### 3.3. Astrocyte subtype composition shifts underlie apparent SLC13A5 stability

Astrocyte supertype proportions changed significantly across pseudoprogression quintiles (Figure 2B). The Astro 2 proportion increased with pseudoprogression (Spearman rho = +0.345, FDR < 0.001), while Astro 3 proportion declined (rho = -0.393, FDR < 0.001). Because Astro 2 carries 24.0% *SLC13A5* prevalence and Astro 3 carries only 0.87%, this compositional shift would be expected to increase overall SLC13A5 prevalence; the expanding subtype expresses the gene at roughly 28-fold higher rates than the contracting one. That the overall prevalence remains stable therefore requires explanation. A concurrent within-Astro 2 transcriptional decline partially offsets the compositional upward pressure: cell-level Spearman correlation between pseudoprogression score and binary *SLC13A5* expression within Astro 2 nuclei yields rho = -0.043 (FDR < 0.05). This effect is small in absolute magnitude and detectable only because of the large n, not because it is biologically large. The net result of these two opposing forces, compositional expansion of the high-expressing subtype and within-subtype transcriptional attenuation, is an apparent stability in overall *SLC13A5* prevalence that conceals a two-layer reorganization of the astrocytic compartment.

### 3.4. Astro 3 is an A1-reactive subtype; Astro 2 is homeostatic

To characterize the functional identities of these two supertypes, we extracted eight canonical reactive astrocyte markers (Figure 3A). Astro 3 showed 6.5-fold higher *C3* prevalence (10.4% vs.∼1.6%; BH-corrected Wilcoxon p < 0.001) and 4.2-fold higher *CD44* prevalence (23.3% vs.∼5.5%; p < 0.001) compared to Astro 2. *C3* and *CD44* are hallmarks of the A1/neurotoxic reactive state induced by microglial IL-1*α*, TNF, and C1q signaling [11]. In contrast, the pan-astrocyte marker *ALDH1L1* was modestly higher in Astro 2 (25.6% vs.∼21.0%; p < 0.001), and *AQP4* prevalence was statistically indistinguishable between subtypes (25.5% vs.∼24.2%; ns), confirming both populations as bona fide astrocytes [7]. Thus, Astro 3 represents an A1-like reactive subtype, while Astro 2 represents a more homeostatic population [12] that is the primary carrier of *SLC13A5*/NaCT expression. Accordingly, donor-level *SLC13A5* prevalence was substantially higher in Astro 2 than in Astro 3 nuclei across all 84 donors (Figure 3B).

**Figure 3.**
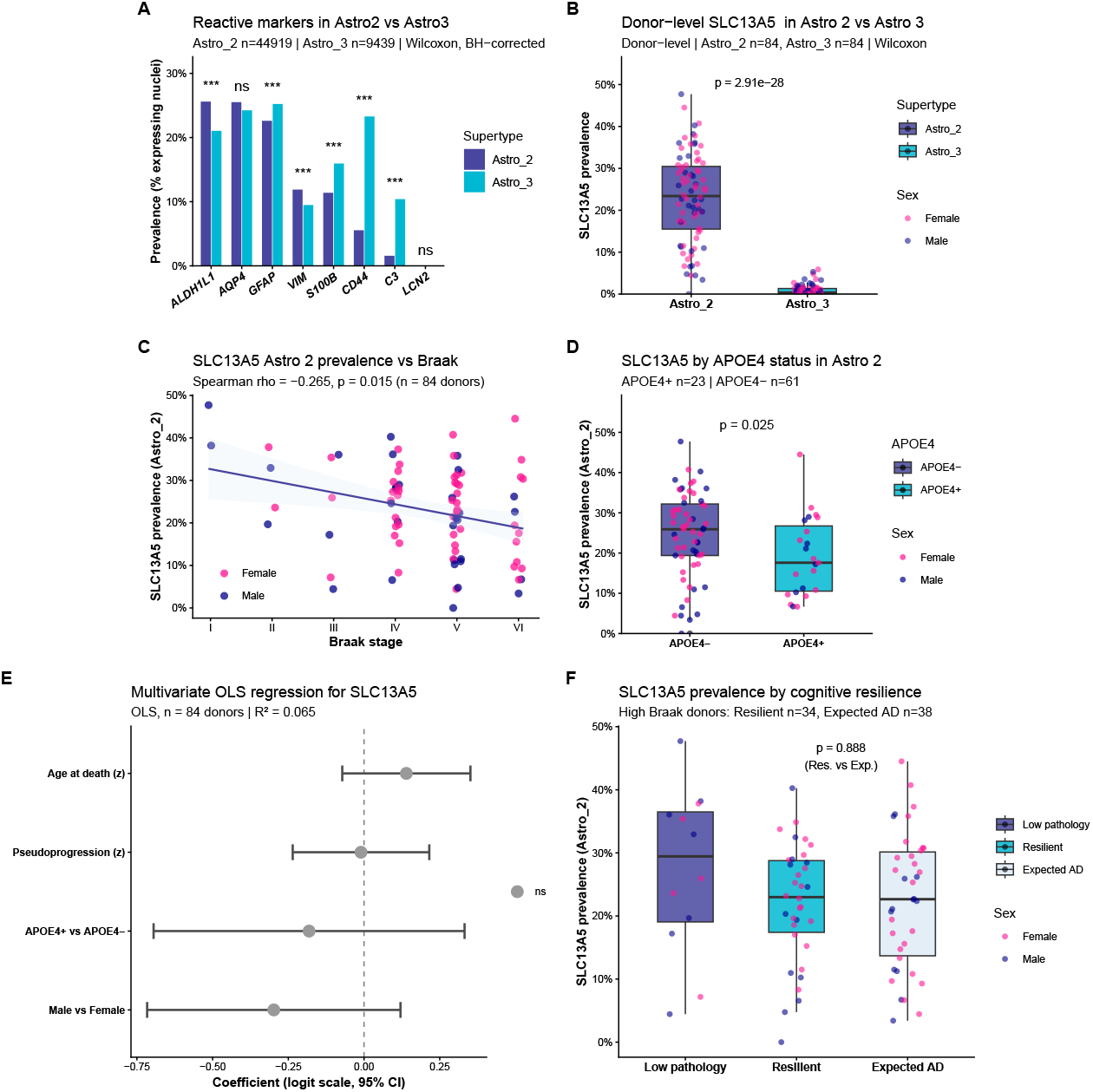
Neuropathological, genetic, and subtype-specific determinants of astrocyte SLC13A5 expression. (A) Reactive astrocyte marker prevalence in Astro 2 versus Astro 3 (Wilcoxon, BH-corrected; *** FDR < 0.001). (B) Donor-level SLC13A5 prevalence in Astro 2 versus Astro 3. (C) SLC13A5 prevalence in Astro 2 nuclei versus Braak stage (Spearman correlation; each point = one donor, colored by sex; regression line with 95% CI). (D) SLC13A5 prevalence in Astro 2 by APOE4 genotype (Wilcoxon rank-sum). (E) Multivariate OLS regression coefficients for predictors of logit-transformed SLC13A5 prevalence (n = 84 donors). (F) SLC13A5 prevalence across cognitive resilience groups (Low pathology: Braak I–III; Resilient: Braak IV–VI + No dementia; Expected AD: Braak IV–VI + Dementia).

### 3.5. Co-expression of citrate metabolism genes in SLC13A5+ nuclei

Phi coefficient analysis revealed weak but positive pairwise co-expression among all eight citrate metabolism genes (phi range: 0.01–0.15). *SLC13A5*+ nuclei showed 2- to 3-fold enrichment for co-expression of *SLC13A3, ACO1*, and *ACLY* compared to *SLC13A5*-nuclei, suggesting that a subset of astrocytes maintains coordinated activity across the citrate import-to-utilization pathway (Figure 2E,F). The highest phi values were observed between metabolic enzymes in the same pathway segment (*SLC25A1*–*ACLY* and *ACO1*–*ACO2*), consistent with shared transcriptional regulation.

### 3.6. SLC13A5 and citrate gene expression correlate with neuropathological burden

At the donor level, *SLC13A5* prevalence in Astro 2 nuclei showed a modest negative association with Braak stage that did not survive BH correction (rho = -0.094, ns; Figure 3C), and negatively correlated with Thal phase (rho = -0.241, p = 0.027) but not with CERAD score (rho = -0.161, ns), suggesting that NaCT expression tracks the progressive spatial spread of amyloid pathology rather than local neuritic plaque density. Thal phase reflects the sequential propagation of amyloid-*β* deposits from allocortical to neocortical regions [16], whereas CERAD quantifies the density of neuritic plaques at the local sampling site [17]; their dissociation here indicates that the regional extent of amyloid spread, rather than its local intensity at the MTG sampling site, is the pathological variable most closely coupled to *SLC13A5* expression in Astro 2.

Sex-stratified analyses revealed that the Braak–SLC13A5 association was driven primarily by male donors (rho = -0.421, p = 0.015, n = 33), while female donors showed no significant correlation (rho = -0.153, p = 0.284, n = 51). A linear interaction model did not detect Braak × sex interaction (*β* = -0.028, p = 0.148), likely reflecting limited power given the male subsample size. This pattern should be interpreted cautiously but warrants investigation in larger cohorts with balanced sex representation.

*SLC13A3* and *ACO1* showed more robust correlations across both pathological measures: *SLC13A3* with Thal (rho = -0.310, BH-adj. p = 0.009) and CERAD (rho = -0.342, p = 0.006); *ACO1* with Thal (rho = -0.307, p = 0.009) and CERAD (rho = -0.349, p = 0.006). Confound checks confirmed that neither age at death (Spearman rho = 0.106, p = 0.337) nor post-mortem interval (rho = 0.119, p = 0.282) meaningfully predicted *SLC13A5* prevalence in Astro 2 nuclei (Figure 3E). Cognitive status (No dementia vs.∼Dementia) was not significantly associated with expression of any of the eight genes after BH correction.

### 3.7. APOE4 genotype is associated with lower SLC13A5 prevalence in Astro 2

APOE4 carriers (n = 23) showed lower median *SLC13A5* prevalence in Astro 2 nuclei compared to APOE4-donors (n = 61; median 17.6% vs.∼25.9%; Wilcoxon p = 0.025; Figure 3D). This difference was not statistically significant across all astrocyte nuclei combined (p = 0.054), consistent with signal dilution by Astro 3 nuclei where *SLC13A5* is nearly absent. In multivariate OLS regression adjusting for pseudoprogression, age, and sex, the APOE4 coefficient did not reach significance (p = 0.484; Figure 3E; R^2^ = 0.065), indicating that the association detected by the non-parametric test is not robust to regression modeling of the skewed outcome distribution. *ACO1* prevalence in Astro 2 showed a similarly borderline APOE4 association (Wilcoxon p = 0.025), while *SLC13A3* and *ACO2* did not differ by APOE4 status.

### 3.8. SLC13A5 expression does not distinguish cognitively resilient from expected-AD donors

To test whether higher NaCT expression might confer cognitive resilience against equivalent tau pathology burden, we compared *SLC13A5* prevalence in Astro 2 between donors with high Braak stage (IV–VI) who retained cognitive function (Resilient; n = 34) versus those with dementia (Expected AD; n = 38). Median Astro 2 *SLC13A5* prevalence was 23.0% in Resilient and 22.7% in Expected AD donors (Wilcoxon p = 0.888; Figure 3F). Kruskal-Wallis test across all three resilience groups (including Low pathology; n = 12) was also non-significant. The cognitive resilience hypothesis for NaCT-mediated citrate transport in MTG astrocytes is not supported by this dataset. These findings are specific to Astro 2 nuclei in the middle temporal gyrus; whether SLC13A5 expression in other brain regions, particularly those where neuronal expression may be more prominent, contributes to cognitive resilience mechanisms cannot be excluded.

## 4. Discussion

This study is the first to characterize *SLC13A5*/NaCT expression dynamics at single-nucleus resolution across the neuropathological continuum of Alzheimer’s disease. Prior snRNA-seq studies of AD have mapped disease-associated transcriptional states in neurons, microglia, and astrocytes [9,10,32,33], but none has examined the citrate transporter gene family or resolved its expression to astrocyte supertypes across a continuous pseudoprogression axis [14]. Three findings stand out from our analysis of 1,378,211 nuclei in the SEA-AD MTG dataset.

Before addressing the three main findings, a foundational observation warrants explicit discussion: SLC13A5 expression in the SEA-AD MTG dataset is nearly absent from neuronal subclasses; the apparent exception, L5 ET neurons (∼7.7% nominal prevalence), was shown to be predominantly attributable to sequencing depth rather than constitutive expression. This is unexpected, as NaCT was originally characterized as a sodium-coupled citrate transporter expressed predominantly in neurons, particularly in the rodent brain [1], and has been described in the literature as a “neuronal citrate transporter.” Its near-absent detection in neuronal nuclei in the human MTG, and its concentration in the homeostatic astrocyte subtype Astro 2, does not necessarily contradict prior work, cell-type expression profiles vary substantially across brain regions, developmental stages, and species, and the MTG may represent a region of astrocyte-dominant SLC13A5 expression independent of what holds in hippocampus, cerebellum, or subcortical structures. Whether Reference donors (neurotypical individuals, underrepresented here) would show detectable neuronal SLC13A5 in this region also remains unresolved. What this dataset establishes clearly is that, in the human MTG across the AD pathological spectrum, the relevant biology of NaCT is astrocytic, and that analyses framed around neuronal function would miss the cell population that carries virtually all detectable expression in this structure.

First, *SLC13A5* expression is highly restricted to astrocytes and concentrated within the Astro 2 homeostatic subtype, which expands as AD pathology progresses while the A1-reactive Astro 3 subtype declines. This compositional shift is the dominant force maintaining overall *SLC13A5* stability. A small but statistically detectable transcriptional decline within Astro 2 (rho = -0.043) is present, though its effect size is near-zero and should not be overinterpreted: it is significant because the cell-level analysis has n = 67,419 observations, not because the effect is biologically large. What matters practically is that NaCT-mediated citrate import is maintained in an expanding homeostatic astrocyte population even as the broader astrocytic compartment undergoes reactive transformation. Compositional restructuring, not transcriptional shutdown, is the operative mechanism.

Second, *SLC13A3* and *ACO1* show more consistent and robust reductions across pseudoprogression, Braak stage, and neuropathological measures of amyloid burden than *SLC13A5* does. The decline in *SLC13A3*, which encodes the Na^+^/dicarboxylate cotransporter NaDC3 responsible for citrate/succinate import into astrocytes [31], has a dual origin: compositional loss of Astro 3 (where *SLC13A3* prevalence is *∼*61%) and transcriptional downregulation within each remaining supertype. *ACO1* has a dual function worth noting: it encodes the cytoplasmic aconitase that interconverts citrate and isocitrate in the TCA shunt, and it acts as iron regulatory protein 1 (IRP1) when its iron-sulfur cluster is disassembled [34]. The observed *ACO1* decline may thus connect to the broader disruption of iron homeostasis reported in AD, independent of its role in citrate metabolism. *SLC13A3* and *ACO1* together may therefore represent more sensitive transcriptional readouts of astrocytic metabolic compromise in AD than *SLC13A5* itself. Their concordant decline across tau (Braak) and amyloid (CERAD, Thal) measures suggests coupling to overall neuropathological burden rather than specificity to a single pathological axis.

Third, the APOE4 association with Astro 2 *SLC13A5* prevalence (p = 0.025 by Wilcoxon) is suggestive but fragile. It fails in multivariate regression (R^2^ = 0.065, all predictors non-significant), and the highly non-normal distribution of the logit-transformed outcome makes the non-parametric test more sensitive to tail differences than the regression model. The grouping used here (APOE4+ defined as E3/E4 and E4/E4, with E2/E4 assigned to APOE4-given the ambiguous risk profile of that genotype) follows the most common convention in AD genetics [20], but a sensitivity analysis including E2/E4 in APOE4+ would be warranted in a larger dataset. The most parsimonious interpretation is a modest distributional shift that may not survive stringent multiple testing. Mechanistically, APOE4 disrupts glial lipid homeostasis through impaired lipid droplet turnover and cholesterol efflux [20,35], and since NaCT activity and membrane trafficking depend on lipid bilayer composition, APOE4-specific lipid dysregulation could reduce effective citrate import capacity. That hypothesis requires testing at the protein level.

Three limitations should be noted. The SEA-AD pseudoprogression score is inferred from transcriptomic signatures rather than measured from neuropathology directly; its relationship to the actual temporal sequence of pathological events is probabilistic. Without spatial information, we cannot determine whether the Astro 2 expansion reflects in-situ transcriptional reprogramming or regional redistribution of astrocyte subtypes across the MTG. And 84 donors, with only 23 APOE4+ among them, leaves the genetic subgroup analyses underpowered.

The finding that cognitively resilient donors show no elevation in Astro 2 *SLC13A5* expression deserves attention. If NaCT-mediated citrate import were a primary mechanism of neuronal metabolic support that buffers against cognitive decline, higher *SLC13A5* expression would be expected in resilient individuals. Its absence is consistent with evidence that cognitive reserve operates through synaptic and network-level factors (education, connectivity, neural efficiency) rather than through individual metabolic gene expression [21]. More broadly, it is important to note that all prevalence estimates in this study reflect mRNA detection in single-nucleus sequencing data. Gene expression does not directly imply protein abundance, subcellular localization, or transport activity, a nucleus counted as SLC13A5-positive carries detectable transcript, but whether the corresponding NaCT protein is present at the plasma membrane and functionally active requires independent validation. Confirmation of NaCT protein expression in human MTG astrocytes, and direct measurement of citrate transport capacity in this cell population, would be needed to translate these transcriptomic findings into functional conclusions. This caveat applies with particular force to the cognitive resilience analysis: if the protective function of NaCT operates primarily at the protein level, it would not be captured by mRNA prevalence in snRNA-seq data, and the null result here would not exclude a functional role.

## 5. Conclusions

Contrary to the canonical description of NaCT as a neuronal citrate transporter, SLC13A5 expression in the human MTG is concentrated in astrocytes, with near-absent detection across neuronal subclasses, a cell-type distribution that reframes the relevant biology of this transporter in this region. Single-nucleus transcriptomics of the SEA-AD MTG dataset shows that *SLC13A5*/NaCT expression is maintained within the expanding Astro 2 homeostatic subtype across Alzheimer’s disease pseudoprogression. The subtype-level dynamics are invisible at the bulk or subclass level and only become apparent when astrocyte populations are resolved individually. The A1-reactive Astro 3 subtype, which contracts with pseudoprogression, carries high *SLC13A3* and *ACO1* expression, and its loss is the dominant source of the observed decline in these metabolic genes across the disease continuum. APOE4 genotype shows a suggestive but statistically fragile association with lower Astro 2 *SLC13A5* expression. Taken together, the data reframe the question of citrate transporter biology in AD from `does NaCT expression change?` to `which astrocyte population changes, and through what mechanism?` Spatial transcriptomics, proteomics, and metabolomics will be needed to determine whether these astrocyte subtype shifts carry functional metabolic consequences.

## Author Contributions

Conceptualization, H.R.F.; methodology, H.R.F.; software, H.R.F.; formal analysis, H.R.F.; data curation, H.R.F.; writing—original draft preparation, H.R.F., P.F.S., G.C.F.; writing—review and editing, H.R.F., P.F.S., G.C.F. The authors have read and agreed to the published version of the manuscript.

## Institutional Review Board Statement

Not applicable. This study used publicly available deidentified data from the Seattle Alzheimer’s Disease Brain Cell Atlas (SEA-AD), available via the Allen Brain Cell Atlas (https://portal.brain-map.org).

## Informed Consent Statement

Not applicable.

## Data Availability Statement

The SEA-AD MTG dataset analyzed in this study is publicly available via the Allen Brain Cell Atlas at https://portal.brain-map.org/atlases-and-data/rnaseq/human-mtg-10x_sea-ad. Analysis code is available from the corresponding author upon reasonable request.

## Acknowledgments

The authors thank the Seattle Alzheimer’s Disease Brain Cell Atlas consortium for making the MTG single-nucleus RNA sequencing dataset publicly available.

## Abbreviations

The following abbreviations are used in this manuscript:

AD: Alzheimer’s disease
snRNA-seq: single-nucleus RNA sequencing
SEA-AD: Seattle Alzheimer’s Disease Brain Cell Atlas
MTG: middle temporal gyrus
NaCT: sodium-coupled citrate transporter
ACLY: ATP-citrate lyase
TCA: tricarboxylic acid
FDR: false discovery rate
OLS: ordinary least squares
LOESS: locally estimated scatterplot smoothing
PMI: post-mortem interval
CERAD: Consortium to Establish a Registry for Alzheimer’s Disease
IRP1: iron regulatory protein 1
UMI: unique molecular identifier

## Disclaimer/Publisher’s Note

The statements, opinions and data contained in all publications are solely those of the individual author(s) and contributor(s) and not of MDPI and/or the editor(s). MDPI and/or the editor(s) disclaim responsibility for any injury to people or property resulting from any ideas, methods, instructions or products referred to in the content.

