## Supplementary material 1 for "Astrocyte subtype-specific expression of the sodium-coupled citrate transporter SLC13A5 and citrate metabolism genes across Alzheimer’s disease pseudoprogression: a single-nucleus RNA sequencing analysis of the human middle temporal gyrus"

### 1. Supplementary Tables

#### *1.1. Table S1. Cell-level Spearman correlations.*

#### *1.2. Table S2. Donor-level Spearman correlations.*

#### *1.3. Table S3. Neuropathological correlations for all eight citrate metabolism genes.*

#### *1.4. Table S4. Reactive astrocyte marker comparison: Astro 2 vs. Astro 3.*

#### *1.5. Table S5. SLC13A5 pseudoprogression correlations per astrocyte supertype.*

#### *1.6. Table S6. Segmented regression of SLC13A5 astrocyte prevalence across pseudoprogression.*

### 2. Supplementary Figures

#### *2.1. Figure S1. Sex-stratified expression trajectories across pseudoprogression.*

#### *2.2. Figure S2. Cognitive status comparison across all eight citrate metabolism genes.*

#### *2.3. Figure S3. SLC13A5 trajectory breakpoint and per-supertype pseudoprogression correlations.*

#### *2.4. Figure S4. Braak stage and APOE4 associations across all eight citrate metabolism genes.*

**Table S1.** Supplementary Table 1. Cell-level Spearman correlations (pseudoprogression vs. binary expression, astrocyte nuclei).

| Gene | Spearman rho | p-value | FDR | Significance | n nuclei |
|---|---|---|---|---|---|
| SLC13A5 | -0.008 | 0.0449 | 0.0609 | ns | 67419 |
| SLC13A3 | -0.080 | 0.0000 | 0.0000 | *** | 67419 |
| SLC25A1 | -0.003 | 0.4890 | 0.4890 | ns | 67419 |
| ACLY | -0.006 | 0.1160 | 0.1330 | ns | 67419 |
| ACO1 | -0.059 | 0.0000 | 0.0000 | *** | 67419 |
| ACO2 | -0.029 | 0.0000 | 0.0000 | *** | 67419 |
| IDH1 | -0.008 | 0.0456 | 0.0609 | ns | 67419 |
| IDH2 | -0.032 | 0.0000 | 0.0000 | *** | 67419 |

**Table S2.** Supplementary Table 2. Donor-level Spearman correlations (mean pseudoprogression vs. bulk astrocyte prevalence, n = 84 donors).

| Gene | Spearman rho | p-value | FDR | Significance | n donors |
|---|---|---|---|---|---|
| SLC13A5 | -0.042 | 0.704000 | 0.78600 | ns | 84 |
| SLC13A3 | -0.362 | 0.000723 | 0.00578 | ** | 84 |
| SLC25A1 | -0.073 | 0.507000 | 0.78600 | ns | 84 |
| ACLY | -0.050 | 0.652000 | 0.78600 | ns | 84 |
| ACO1 | -0.322 | 0.002770 | 0.01110 | * | 84 |
| ACO2 | -0.258 | 0.017700 | 0.04710 | * | 84 |
| IDH1 | -0.030 | 0.786000 | 0.78600 | ns | 84 |
| IDH2 | -0.170 | 0.122000 | 0.24500 | ns | 84 |

**Table S3.** Supplementary Table 3. Neuropathological correlations for citrate gene prevalences. Braak correlations use donor-level prevalence across all astrocytes (n = 84 donors). Thal and CERAD correlations are Astro 2-specific (SLC13A5, SLC13A3, ACO1, ACO2 only).

| Gene | Pathology measure | Spearman rho | p-value | FDR | Sig. | n donors |
|---|---|---|---|---|---|---|
| ACLY | Braak stage | -0.086 | 4.38e-01 | 0.438000 | ns | 84 |
| ACO1 | Braak stage | -0.430 | 4.51e-05 | 0.000361 | *** | 84 |
| ACO1 | CERAD score | -0.349 | 1.13e-03 | 0.005760 | ** | 84 |
| ACO1 | Thal | -0.307 | 4.49e-03 | 0.008970 | ** | 84 |
| ACO2 | Braak stage | -0.198 | 7.05e-02 | 0.113000 | ns | 84 |
| ACO2 | CERAD score | -0.076 | 4.92e-01 | 0.492000 | ns | 84 |
| ACO2 | Thal | -0.156 | 1.57e-01 | 0.179000 | ns | 84 |
| IDH1 | Braak stage | -0.212 | 5.29e-02 | 0.106000 | ns | 84 |
| IDH2 | Braak stage | -0.310 | 4.17e-03 | 0.016100 | * | 84 |
| SLC13A3 | Braak stage | -0.297 | 6.04e-03 | 0.016100 | * | 84 |
| SLC13A3 | CERAD score | -0.342 | 1.44e-03 | 0.005760 | ** | 84 |
| SLC13A3 | Thal | -0.310 | 4.14e-03 | 0.008970 | ** | 84 |
| SLC13A5 | Braak stage | -0.094 | 3.95e-01 | 0.438000 | ns | 84 |
| SLC13A5 | CERAD score | -0.161 | 1.42e-01 | 0.179000 | ns | 84 |
| SLC13A5 | Thal | -0.241 | 2.73e-02 | 0.043700 | * | 84 |
| SLC25A1 | Braak stage | -0.091 | 4.08e-01 | 0.438000 | ns | 84 |

https://doi.org/10.3390/cimb1010000

**Table S4.** Supplementary Table 4. Wilcoxon rank-sum comparison of reactive astrocyte marker prevalences between Astro 2 and Astro 3 supertypes (astrocyte nuclei, BH-corrected).

| Marker | Prevalence Astro 2 | Prevalence Astro 3 | log2FC (A2/A3) | p-value | FDR | Sig. |
|---|---|---|---|---|---|---|
| GFAP | 0.226 | 0.252 | -0.16 | 0.000 | 0.000 | *** |
| VIM | 0.119 | 0.095 | 0.33 | 0.000 | 0.000 | *** |
| C3 | 0.016 | 0.104 | -2.74 | 0.000 | 0.000 | *** |
| ALDH1L1 | 0.256 | 0.210 | 0.28 | 0.000 | 0.000 | *** |
| AQP4 | 0.255 | 0.242 | 0.07 | 0.363 | 0.415 | ns |
| S100B | 0.114 | 0.159 | -0.49 | 0.000 | 0.000 | *** |
| CD44 | 0.055 | 0.233 | -2.07 | 0.000 | 0.000 | *** |
| LCN2 | 0.000 | 0.000 | Inf | 0.517 | 0.517 | ns |

**Table S5.** Supplementary Table 5. SLC13A5 Spearman correlations with pseudoprogression per astrocyte supertype. Prevalence: binary expression vs. pseudo-score across all nuclei in each supertype (Astro 4 excluded; < 50 expressing cells). Intensity: log1p-transformed counts among SLC13A5-expressing nuclei only (Astro 3 and Astro 4 additionally excluded due to < 100 expressing cells).

| Supertype | Measure | n nuclei | n expressing | Spearman rho | p-value | FDR |
|---|---|---|---|---|---|---|
| Astro_1 | Intensity (SLC13A5+ only) | 399 | 399 | -0.004 | 9.30e-01 | 9.30e-01 |
| Astro_1 | Prevalence (binary) | 5335 | 399 | -0.059 | 1.55e-05 | 3.87e-05 |
| Astro_2 | Intensity (SLC13A5+ only) | 10800 | 10800 | 0.042 | 1.08e-05 | 4.31e-05 |
| Astro_2 | Prevalence (binary) | 44919 | 10800 | -0.043 | 0.00e+00 | 0.00e+00 |
| Astro_3 | Prevalence (binary) | 9439 | 82 | 0.015 | 1.34e-01 | 1.34e-01 |
| Astro_5 | Intensity (SLC13A5+ only) | 452 | 452 | -0.055 | 2.45e-01 | 3.26e-01 |
| Astro_5 | Prevalence (binary) | 3603 | 452 | -0.037 | 2.83e-02 | 3.54e-02 |
| Astro_6 | Intensity (SLC13A5+ only) | 732 | 732 | 0.046 | 2.10e-01 | 3.26e-01 |
| Astro_6 | Prevalence (binary) | 3271 | 732 | 0.043 | 1.29e-02 | 2.15e-02 |

**Table S6.** Supplementary Table 6. Segmented ordinary least-squares regression of SLC13A5 astrocyte prevalence across 20 equal-width pseudoprogression bins (n = 19 bins with data; weighted by nuclei per bin). Davies test evaluates whether the breakpoint improves fit over a linear model.

| Parameter | Value |
|---|---|
| Breakpoint (pseudo-score) | 0.5195 |
| Breakpoint SE | 0.0992 |
| Slope before breakpoint (per unit pseudo-score) | 0.3319 |
| Slope after breakpoint | -0.1940 |
| Adjusted $R^2$ | 0.4143 |
| Davies test p-value (breakpoint significance) | 0.0392 |

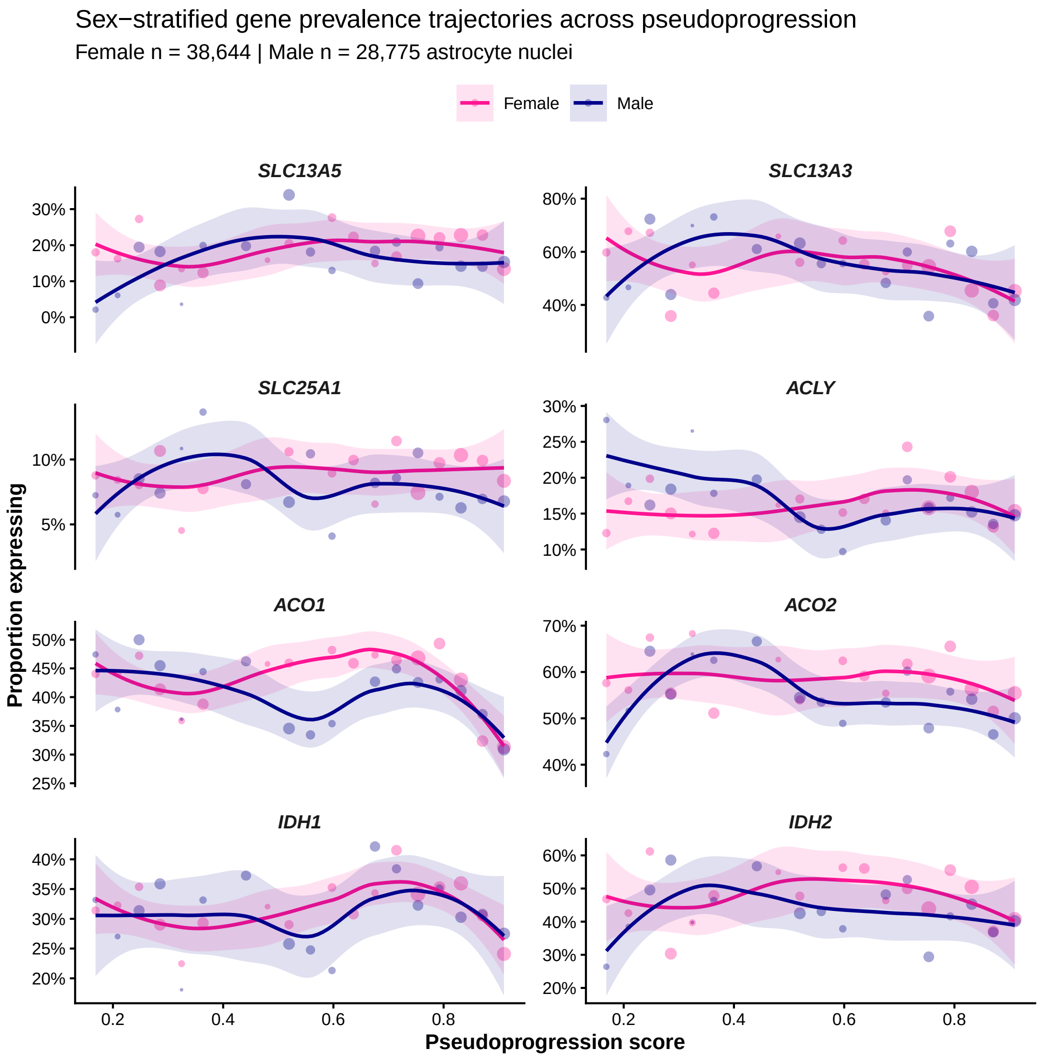


**Figure S1.** Figure S1. Sex-stratified LOESS-smoothed prevalence trajectories for all eight citrate metabolism genes across the pseudoprogression score (20 equal-width bins; shading = 95% CI). Each point represents one pseudoprogression bin; point size reflects the number of astrocyte nuclei in that bin.

https://doi.org/10.3390/cimb1010000

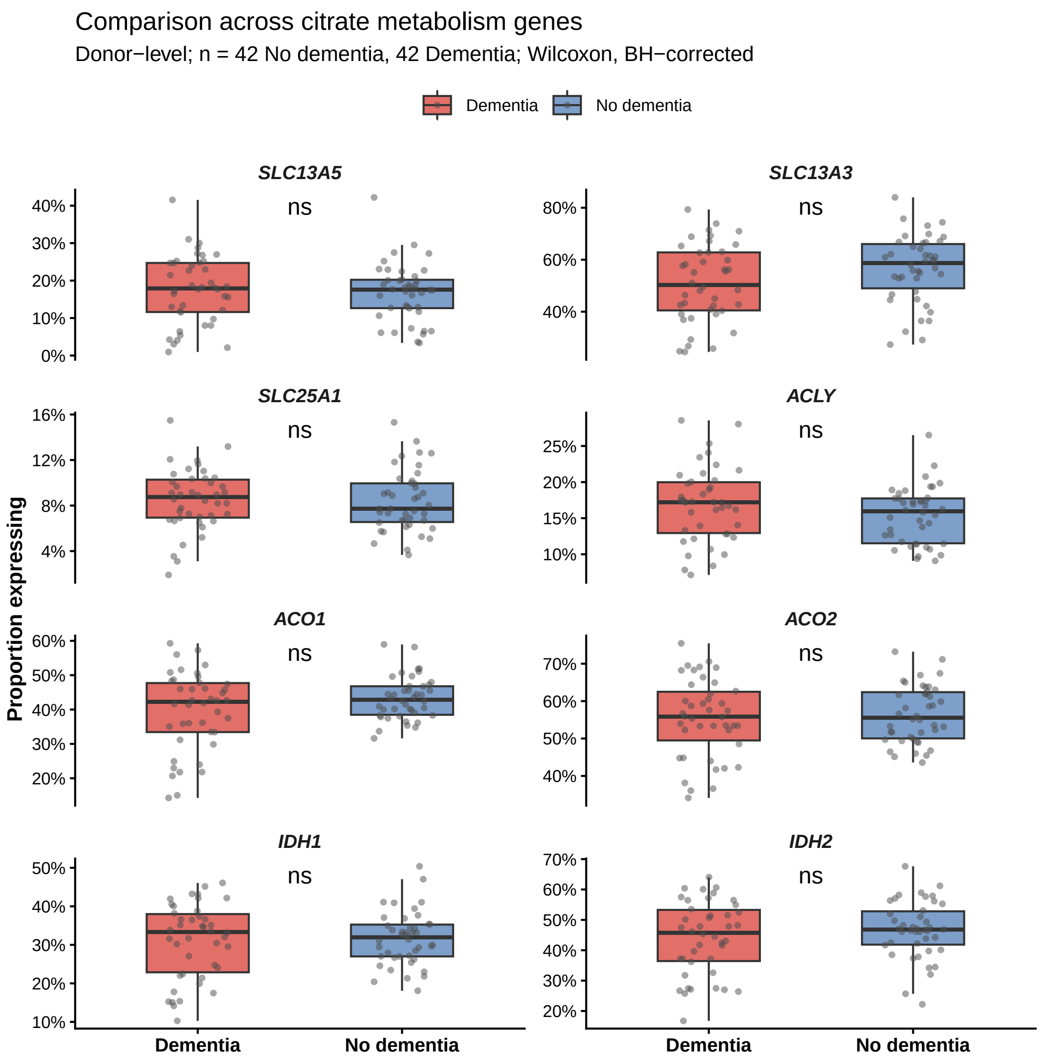


**Figure S2.** Figure S2. Donor-level prevalence of each citrate metabolism gene stratified by cognitive status (No dementia vs. Dementia; Reference donors excluded). Boxes show median and IQR; significance labels reflect BH-corrected Wilcoxon rank-sum tests (*** FDR < 0.001, ** FDR < 0.01, * FDR < 0.05, ns = not significant).

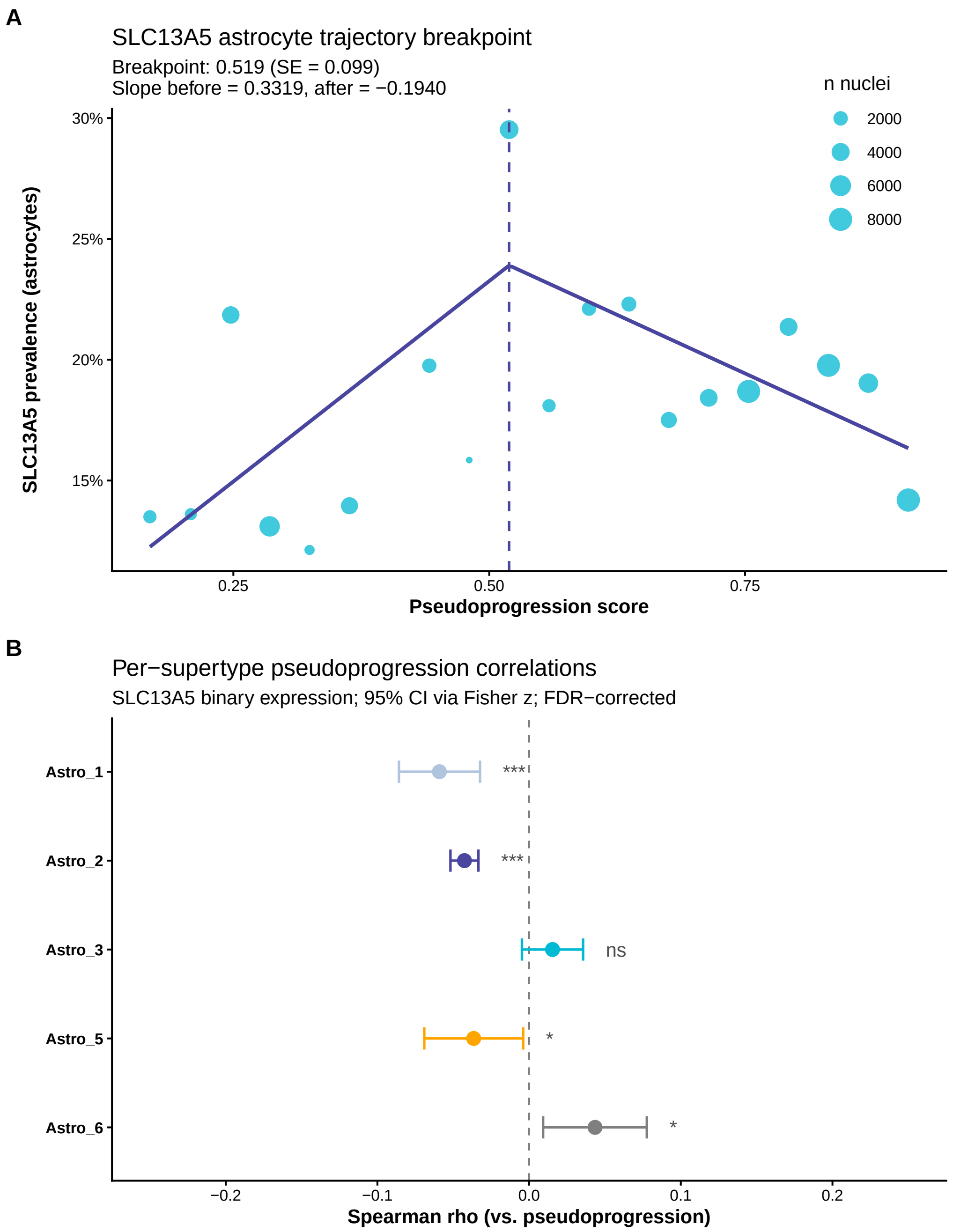


**Figure S3.** Figure S3. (**A**) Segmented OLS regression of SLC13A5 prevalence across 20 pseudoprogression bins in astrocytes. The dashed vertical line marks the estimated breakpoint; the purple line is the piecewise-linear fit weighted by nuclei per bin. (**B**) Per-supertype Spearman correlations of SLC13A5 binary expression with pseudoprogression score. Error bars = 95% CI via Fisher z-transformation. FDR-corrected significance labels (*** < 0.001, ** < 0.01, * < 0.05, ns).

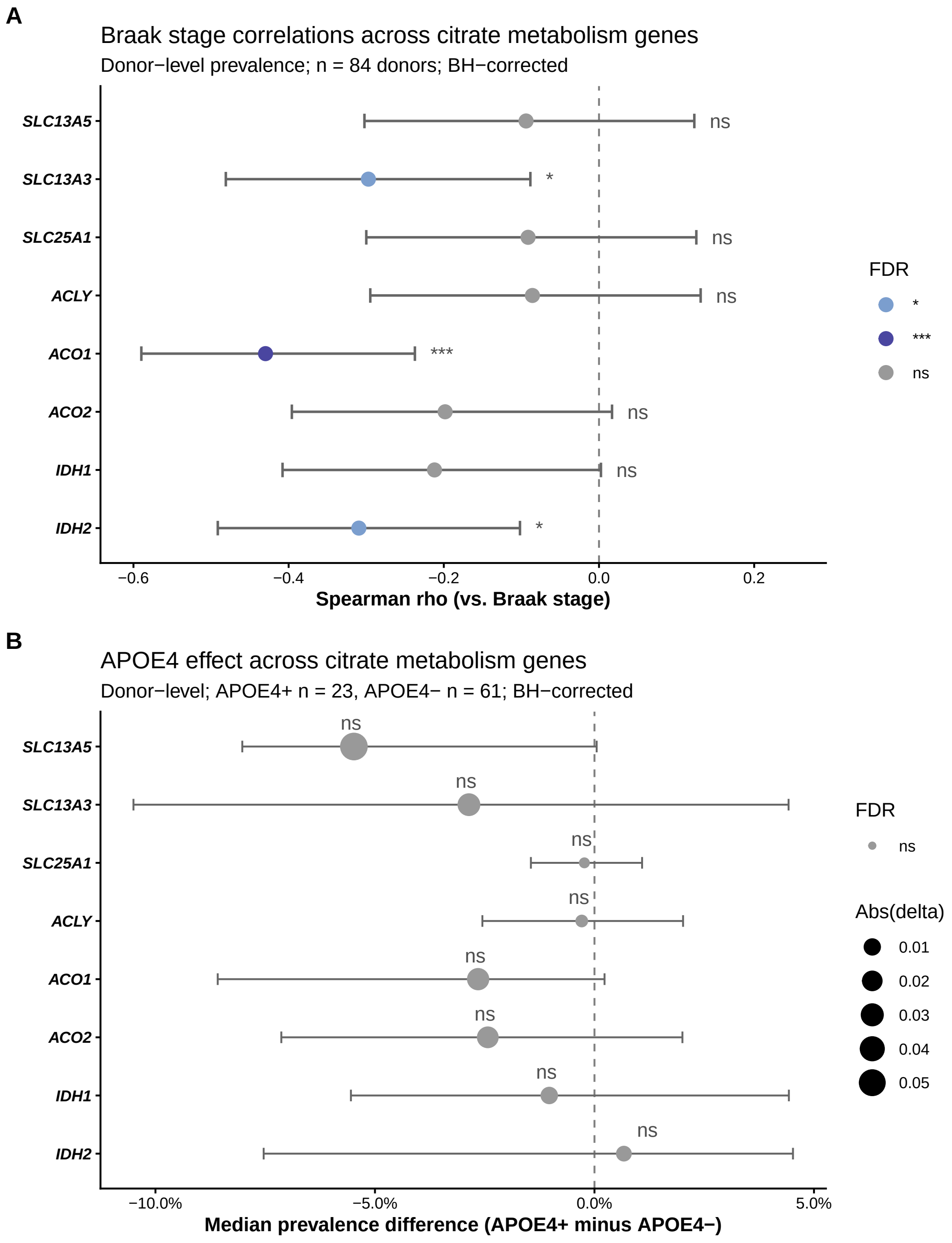


**Figure S4.** Figure S4. (**A**) Donor-level Spearman correlations between Braak neurofibrillary tangle stage and bulk astrocyte prevalence for each gene (n = 84 donors; 95% CI via Fisher z-transformation; BH-corrected). (**B**) Difference in median donor-level prevalence between APOE4 carriers and non-carriers (Wilcoxon rank-sum; BH-corrected). Point size is proportional to the absolute difference.
